# *In-Silico* Investigation Reveals a Potential Functional Role of Human Microbiome in Chronic Obstructive Pulmonary Disease

**DOI:** 10.1101/2025.06.29.660899

**Authors:** Nilesh Jana, Oindrila Dhara, Sukanta S. Bhattacharya

**Affiliations:** Department of Biosciences, JIS University, Kolkata

**Keywords:** Chronic Obstructive Pulmonary Disease, Inflammation, Microbiome

## Abstract

Chronic Obstructive Pulmonary Disease (COPD) is a progressive enervating lung disease characterized by chronic inflammation, airway inhibition and unrecoverable structural damage to the lungs. While traditionally associated with environmental factors similar as cigarette smoke and air pollution as well as genetic factors, recent revelations has increasingly indicative of the role of microbiomes in the modulation of this disease. This study explores the structural and functional relations between microbial proteins and human proteins involved in COPD using in-silico bioinformatic tools. Several critical human proteins involved in COPD pathogenesis such as MMP 1, MMP 2, MMP 14, TNF-α, TGF-β were evaluated for the presence of microbial homologous proteins using BLAST analysis in the microbial database. The results revealed significant structural homology with microbial proteins from species including *E. coli, K.pneumonia, Bacillus thuringiensis, Bifidobacterium samirii*. This microbial protein may impact COPD progression through mechanisms similar as molecular belittlement and modulation of host vulnerable responses. Further molecular docking simulations were conducted using the herbal drug oleanolic acid known for their anti-inflammatory effect and can be potentially used in the treatment of COPD. Results demonstrated favourable interactions between these composites and both human and microbial proteins, indicating therapeutic potential of the compounds. For case oleanolic acid showed strong interaction with TNF-α and its microbial homolog. Also phylogenetic tree construction using Clustal Omega handed perceptivity into the evolutionary connections between the bacteria hosting homologous proteins, suggesting possible ancestral links and participated functional pathway applicable to COPD pathology. The finding emphasize the possibility of microbiome deduced proteins to act as modifiers in COPD progression as well as the therapeutic potential of the herbal compounds in targeting both host and microbial factors, which need to be further ascertained by in vitro, in vivo, as well as clinical studies. This novel in silico approach offers a unique prospective on COPD treatment by integrating microbiome and phytotherapy laying the groundwork for future personalized therapeutic strategies targeting both microbial and host inflammatory pathways.

## 1. Introduction

Chronic Obstructive Pulmonary Disease (COPD) is the third most common cause of death by 2020 and is currently one of the main causes of morbidity and mortality globally. COPD is the fourth most common causes of death in the US, and its prevalence and death rates are continually rising. Chronic obstructive pulmonary disease (COPD) is a lung disease that results in restricted airflow and breathing difficulties (Thawanaphong and Nair 2025; Mi et al. 2025). This is a lung condition that worsens with time and is not easily cured. It is occasionally referred to as chronic bronchitis or emphysema. People with COPD may feel lung damage or excessive mucus secretion. Symptoms are fatigue, wheezing, breathing difficulties, and coughing, occasionally with excessive mucus secretion(MacLeod et al. 2021; Huang et al. 2025). The most prevalent causes of COPD are air pollution and smoking. Epidemiological data suggests that COPD is also triggered by environmental and occupational exposures which exposes the subjects to chemicals, toxicants such as pesticide, cleaning supplies among others. Occupational risk for COPD is also increased by long term exposure to vapours, gases, dust and fumes in sectors such as manufacturing, building, mining, and agriculture. Specially impacted are occupations like plastic molders, food processors, painters, gardeners. Prevention and awareness campaigns are crucial to lowering the number of occupational COPD cases (De Matteis 2022).

Several other factors such as the genetic features, inflammatory responses, microbiome, oxidative stress may also play a critical role. Genetic factors such as gene polymorphisms are important in the development of COPD. Increased vulnerability to the disease has been associated with mutations in matrix metalloproteinases (MMPs), tumor necrosis factors (TNF-α), Alpha 1 antitrypsin (AAT). Among the inflammatory responses, chronic inflammation involving monocytes, neutrophils, and CD8+ lymphocytes is the hallmark of COPD, which results in long lasting airway damage. High levels of proinflammatory cytokines such as TNF-α and Interleukin-8 (IL-8), are present in COPD patients. Recent findings are also indicative of a role of human microbiome in the onset of the disease. Prolonged bacterial infections, especially those caused by normal microflora (*Haemophilus influenza*) can cause the lung to become inflamed. By activating lung T cells and raising reactive oxygen species and protease activity, these infections worsen the symptoms of COPD. Also, the microbial infections environmental pollutants both contribute to oxidative stress, which exacerbates tissue damage and lung inflammation (Ritchie and Wedzicha 2020). The lung microbiome especially in the lower airway, has been linked to disease progression. The upper and lower airways have separate microbial communities, and their density diminishes as the respiratory system descends, Bacterial or viral infections are responsible for 50% -60% of COPD exacerbations, leading to increased mucus production and decreased lung fuction (Suyun Yu 2023). In order to help patients with COPD avoid acute respiratory tract infections, it is critical to strengthen their immune systems. The following bacteria produce bacterial lysates through alkaline proteolysis: *Staphylococcus aureus, Steptococcus pyogenes, Klebsiella pneumonia, Haemophilus influenzae, E coli*. Bacterial lysates have been demonstrated to have pleotropic immunomodulatory effect on both the innate and adaptive immune responses in addition to eliminating pathogens (Yafei Qi 2022). It has been recently reported that the gut condition also play a role in COPD progression. Dysbiosis of the intestinal microenvironment effects importantly to the establishment of COPD. Intestinal microflora produces organic acids that down regulate total cell and macrophage count, lymphocyte and neutrophil counts in BALF that results in anti-inflammatory effects in lungs-suppress elastase induced emphysema (Zhi Song 2024).

According to (Yu et al. 2023) pathway of COPD involves : Microbiome changes in COPD, Inflammation and microbial dysbiosis, Microbial metabolites function, The COPD gut lung axis, Microbiota changes and COPD exacerbation, Microbiome’s modulation therapeutic potential. these pathway involves microbiome. The microbial diversity in the lungs of people with COPD is lower than that of healthy people. Pathogenic bacteria (*H. influenzae, Pseudomonas aeruginosa*) take the lead as the proportion of Proteobacteria to Firmicutes rises. Chronic inflammation is brought on by disturbances in the lung microbiota. Lung tissue damage is exacerbated by dysbiosis which causes an increase in the secretion of pro inflammatory cytokines (IL 1 Beta, IL 6, TNF alpha).

By affecting immunological responses and inflammation, the lung microbiota interacts with COPD; dysbiosis accelerates the course of the disease by impairing immune control and increasing airway inflammation. Exacerbations, worse lung function and higher mortality are linked to changes in the microbiota such as rise in Proteobacteria and a fall in Firmicutes (Russo et al. 2022). By affecting immunological responses and inflammation, the microbiome in COPD contributes significantly to the course of the disease and its flare ups. The severity of the condition affects the diversity of bacteria in the lungs and the presence of the pathogenic bacteria can lead to acute exacerbations, airway remodelling and chronic inflammation (Martinez et al. 2013). Because changes in gut microbiota, specially a decline in Bacteroidetes, results in less generation of beneficial short chain fatty acids (SCFAs), which help control inflammation, the gut lung axis is important in COPD. This imbalance exacerbates the course of COPD by worsening lung function and contributing to systemic inflammation (Cheng and Zhang 2024).

However, despite the microbiome dysbiosis observed in the patients as well as smokers and people exposed to toxicants which are linked to COPD, a clear mechanistic or functional role of the microbiome is yet to be established. In the present report we investigated the structural homology between microbial and human proteins as well as evaluated the interactions of two herbal drugs, namely oleanoic acid and methanol, with the proteins of our interest using in-silico tools. To the best of our knowledge, this is the first report on the structural homology between the microbial proteins and human proteins playing a role in the progression of COPD.

## 2. Materials & Methods

### 2.1 Pathway Evaluation

An assessment of the factors playing a role in COPD inflammation and fibrosis pathway was carried out, encompassing important mediators such as IL-13, Connective Tissue Growth Factor (CTGF), Tumor Necrosis Fibrosis alpha (TNF-α), Transforming Growth Factor beta (TGF-β), Matrix metalloproteinases (MMPs) and Monocyte Chemoattractant Proteins (MCPs). For the purpose, a thorough literature review was carried out to assess how these molecular markers contribute to the development of COPD through airway remodelling, extracellular matrix breakdown and chronic inflammation. Furthermore, pathway enrichment analysis based on bioinformatics was carried out to confirm the role of these mediators in the pathophysiology of COPD. The literature review’s conclusions, which demonstrated the biomarker’s importance in COPD related inflammation and fibrosis, served as a guide for the selection of these indicators.

### 2.2 Assessment of homology of human proteins and microbial proteins

NCBI database was used to obtain the FASTA sequence of the targeted human proteins. A BLAST analysis was carried out using the obtained FASTA sequence against the microbial database. From the description table, the microbial proteins with highest percentage identity (Homology) protein for further investigation.

### 2.3 Evaluation Of Protein Drug Interaction

For this at first selection of the proteins and drugs was done which will be used for molecular docking. Firstly search the desired proteins in NCBI and the FASTA sequences was copied. Using the online modelling tool (https://swissmodel.expasy.org/ accessed on May 15^th^, 2025) the protein model with the highest percentage identity was created and downloaded in PDB format using standard protocol. For obtaining the 3D structure of the drug molecule to be used, PubChem database was searched with the name of the desired drug and 3D structure downloaded in SDF format. Openbabel software was open and the SDF drug file was converted to pdbqt format. Autodock 4 option from Amdock was then selected receptor was selected as the PDB protein file and ligand was selected as the pdbqt drug file and Prepare input option was clicked, wait for completion, then “Define search space” option was clicked and wait for completion. After that Run option was clicked and wait for the completion. After completion of the task, it was visualized, and analysis carried out using PyMOL.

### 2.4 Softwares and Online tools

Amdock and PyMOL softwares were used for 3D protein structures and studying the interactions while Openbabel software was used to determine the 3D structure of the ligand.

## 3. Results and Discussion

Several microbial proteins were found to share structural homology with the selected human proteins (Table 1). In the table given below, the microbial protein sharing the structural homology with human protein has been detailed along with the microorganism (Bacteria) producing the protein. The proteins showing a homology more than 50% were taken for further analysis using docking studies. In the results representing the docking (Figures 1 to 4), the left column shows the human proteins 3D structure and its interactions with the drug molecule while the right column shows the same about the microbial proteins (Homologous to human protein) as visualized by PyMOL. All the above proteins are docked with oleanolic acid, a herbal drug which is derived from Olive tree (*Olea europaea*). It have anti-inflammatory effect, antioxidant, antifibrotic properties. So it can be used for management of the COPD disease though the disease is not completely cure by using this compound but the severity of the disease can be low. The data in the figures also shows the details of the binding site, affinity (kcal/mol), estimated Ki, KI units, ligand efficiency. More negative binding affinity value (kcal/mol) in a docking table indicates a stronger and stable binding interactions between a ligand and its target. We evaluated the interaction of oleanolic acid (green) with TNF-α (Cyan), which can be visualized as the bound structure within a helical region of the protein using molecular docking (Fig. 1). Binding site 4 demonstrates the highest affinity (-6.3 kcal/mol) and the lowest estimated Ki (24.10 uM) indicates it as the most favorable binding site for oleanolic acid on TNF-α (Fig. 1(c)). The molecular docking interaction of oleanolic acid (green) with EF Hand domain protein of the myosin regulatory light chain (Cyan), positioned in a helical region of the protein structure (Fig. 1(d)). It shows the binding site 3 of EF Hand Domain Protein has the highest binding affinity (-7.91 kcal/mol) and the lowest estimated Ki (1.59 uM) indicates it is the most favorable site for oleanolic acid. Molecular docking of oleanolic acid within the TGF-β receptor binding pocket (Fig. 2(c)) were also carried out, which indicated stable interactions through hydrogen bonds and hydrophobic contacts. It quantified the binding affinity, with binding site 9 showing the strongest interaction (−7.67 kcal/mol, Ki = 2.39 uM), suggesting it as the most favorable site for ligand binding. The docking conformation of oleanolic acid within the active site of the kinase protein, showing tight binding supported by multiple interactions were also visualized (Figure 2(d)). It reveals strong binding affinity at site 3 (−9.79 kcal/mol, Ki = 66.66 nM), indicating oleanolic acid has high nanomolar potency and efficient ligand binding to kinase protein. 3D molecular docking pose of oleanolic acid bound within the active site of MMP 14, indicating proper orientation and interaction within the binding pocket (Fig. 3(c)). It revealed strong binding affinity at site 3 (−9.79 kcal/mol, Ki = 66.66 nM), indicating oleanolic acid has high nanomolar potency and efficient ligand binding to MMP 14. (Fig 3(d)) the 3D molecular docking of oleanolic acid with apolipoprotein, demonstrating a stable binding conformation within the helical structure. This interaction between apolipoprotein with oleanolic acid where the best binding site exhibits a moderate binding affinity of -7.55 kcal/mol and a ligand efficiency of -0.23. Docking interactions between oleanolic acid and MMP-2 are visualized with oleanolic acid (green) is bound to the active site of Human MMP-2 protein (cyan), indicating specific interactions like hydrogen bonds (Fig. 4(c)). Binding site 2 shows the strongest interaction with a binding affinity of -8.54 kcal/mol and an estimated Ki of 549.72 nM, suggesting favorable binding of oleanolic acid to MMP-2. The interaction of the microbial homolog of MMP-2, that is Sugar Porter Family MFS transporter with oleanolic acid, is visualized with the oleanolic acid bound within the active site of the Sugar Porter family MFS transporter, indicating stable binding via key interactions (Fig 4(d)). It shows binding Site 5 with the best binding affinity of -9.27 kcal/mol and a low estimated Ki of 160.34 nM, suggesting high binding potency of oleanolic acid at this site.. Oleanolic acid is a type of herbal drug which is found in olive tree. It shows effect in treatment for COPD due to its anti-inflammatory, antioxidant, activity. Reports indicate that it can reduce inflammation in the lungs, improve lung function, and protect against lung damage (Bayülgen et al. 2019).

**Figure 1:**
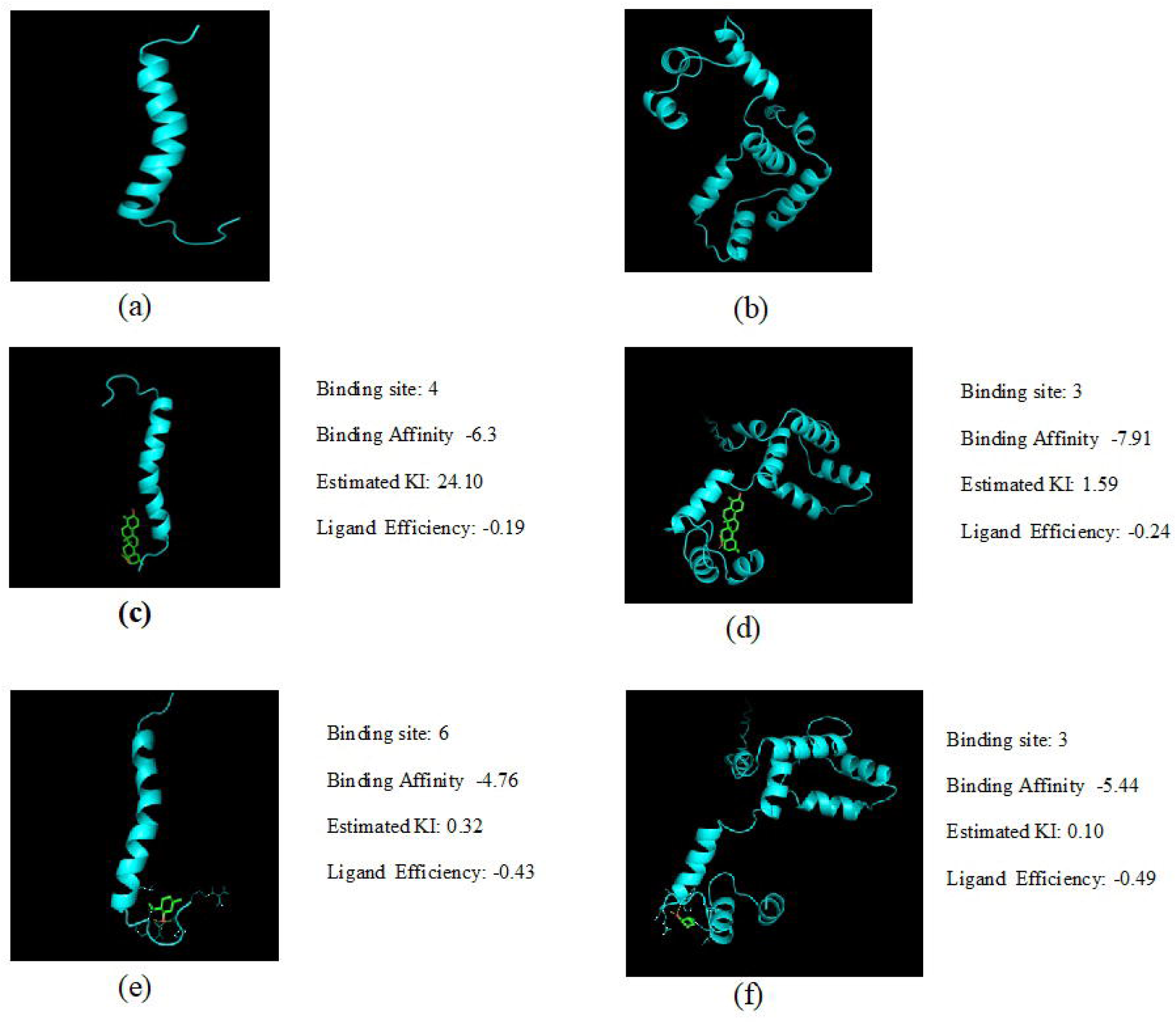
(a) Human TNF Alpha protein (b) Microbial homologue (EF Hand Domain protein of Myosin regulatory Light Chain); Docking result (3D visualization) of (c) TNF Alpha with Oleanolic acid (d) EF Hand Domain protein of Myosin regulatory Light Chain with Oleanolic acid (e) TNF Alpha with Menthol (f) EF Hand Domain protein of Myosin regulatory Light Chain with Menthol

**Figure 2:**
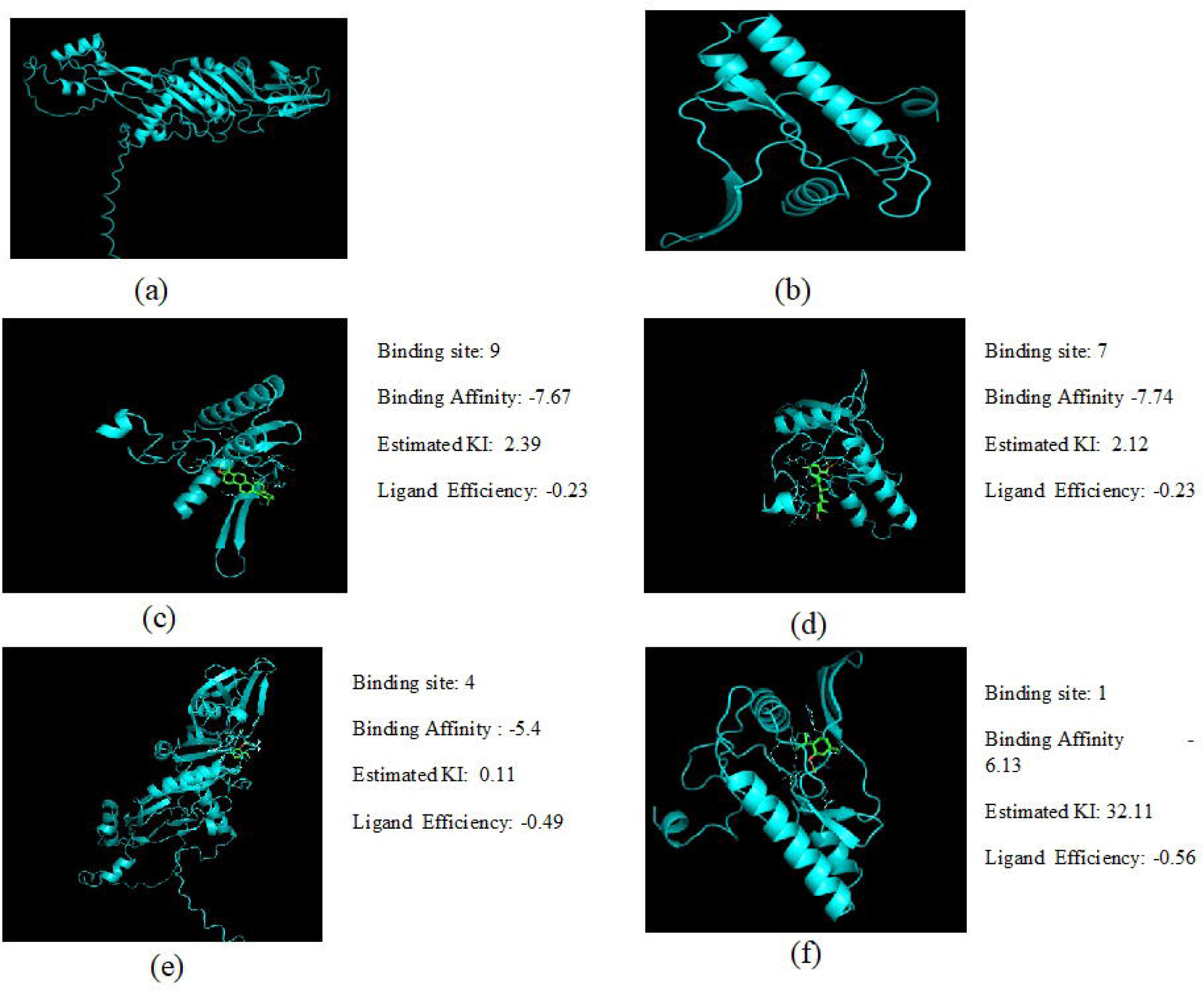
(a) Human TGF Beta protein (b) Microbial homolog Protein kinase; Docking result (3D visualization) of (c) TGF Beta with Oleanolic acid, (d) Protein kinase with Oleanolic acid, (e) TGF Beta with Menthol, (f) Protein kinase with Menthol

**Figure 3:**
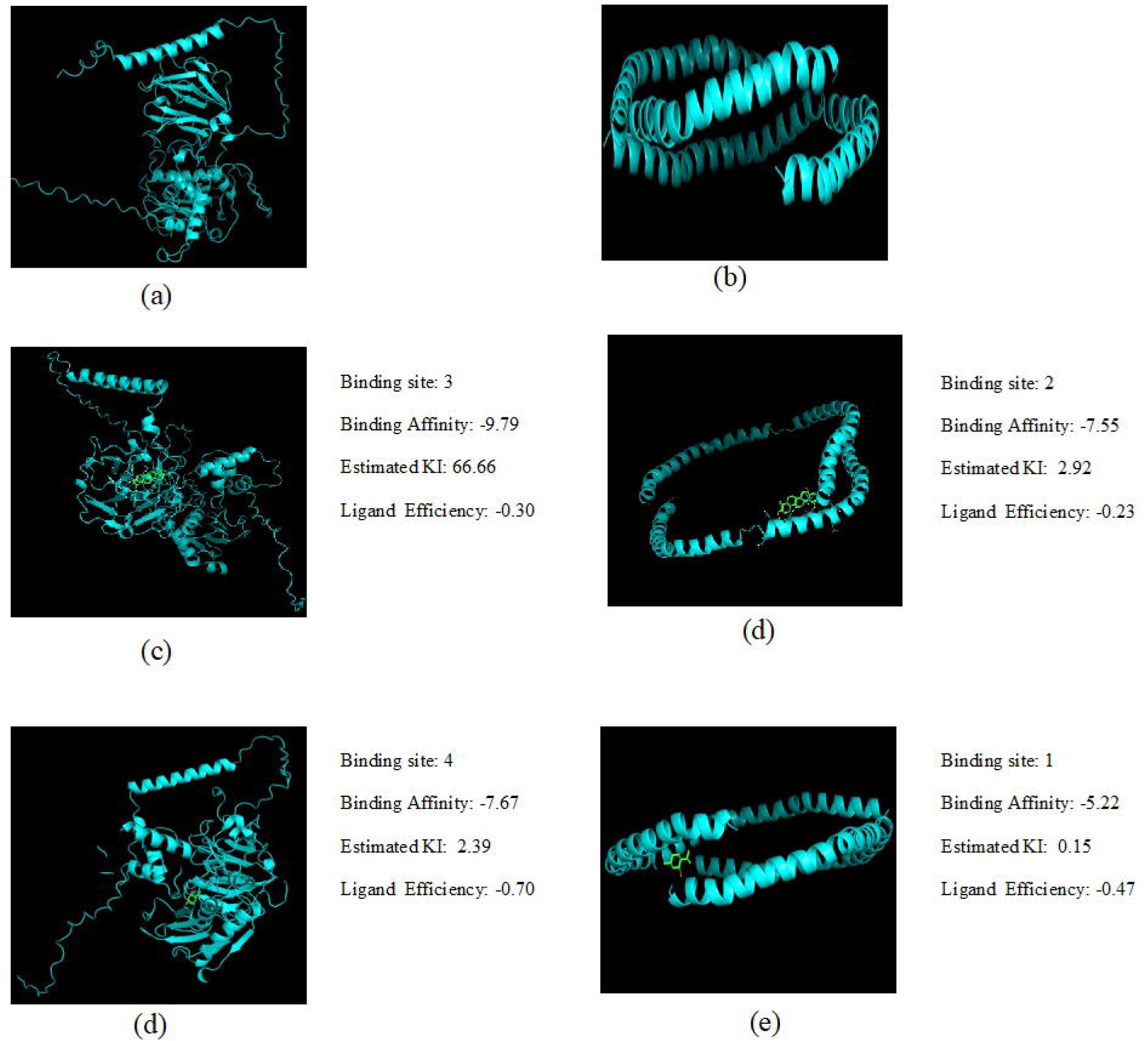
(a) Human MMP 14 protein Fig 5.15: Microbial Apolipoprotein A1/A4 homologue Fig 5.16: Docking result (3D visualization) of MMP 14 with Oleanolic acid Fig 5.17: Docking result (3D visualization) of Apolipoprotein A1/A4 with Oleanolic acid Fig 5.18: Docking result (3D visualization) of MMP 14 with Menthol Fig 5.19: Docking result (3D visualization) of Apolipoprotein A1/A4 with Menthol

**Figure 4:**
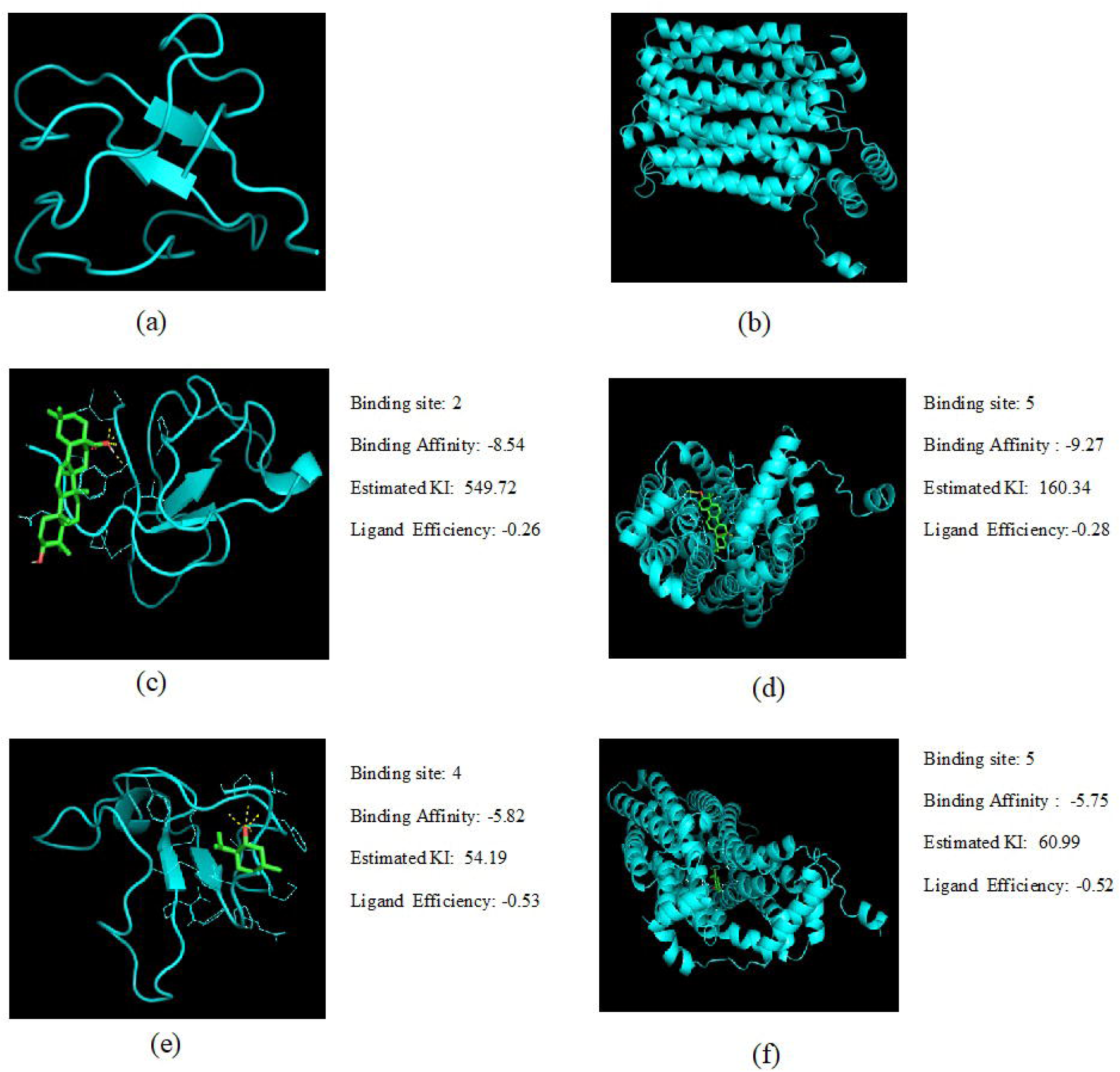
(a) Human MMP 2 protein (b) Microbial Sugar porter family MFS transporter homologue; Docking result (3D visualization) of (c) MMP 2 with Oleanolic Acid (d) Sugar porter family MFS transporter with Oleanolic Acid (e)MMP 2 with Menthol (f)Sugar porter family MFS transporter with Menthol

**Table 1:**
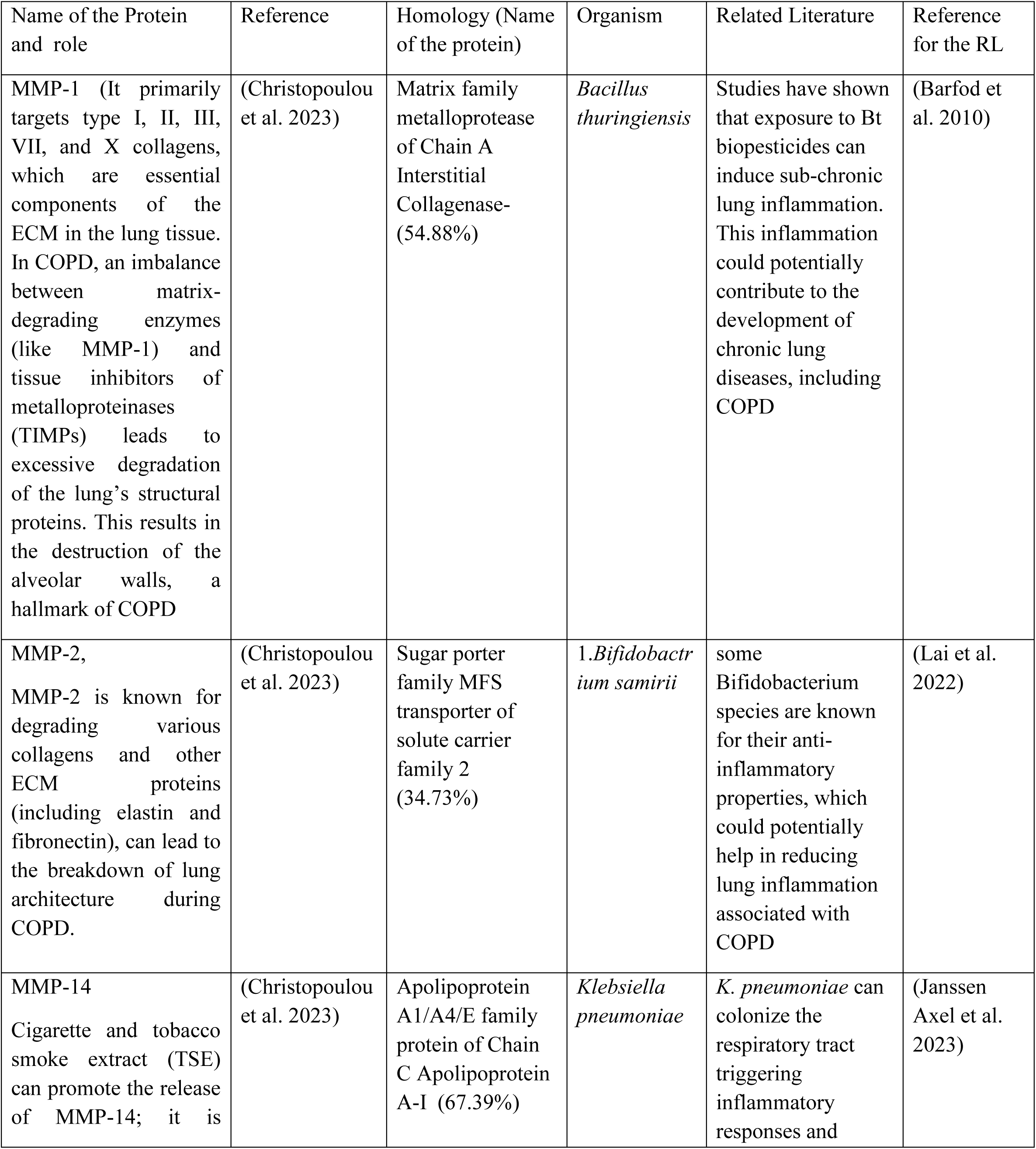
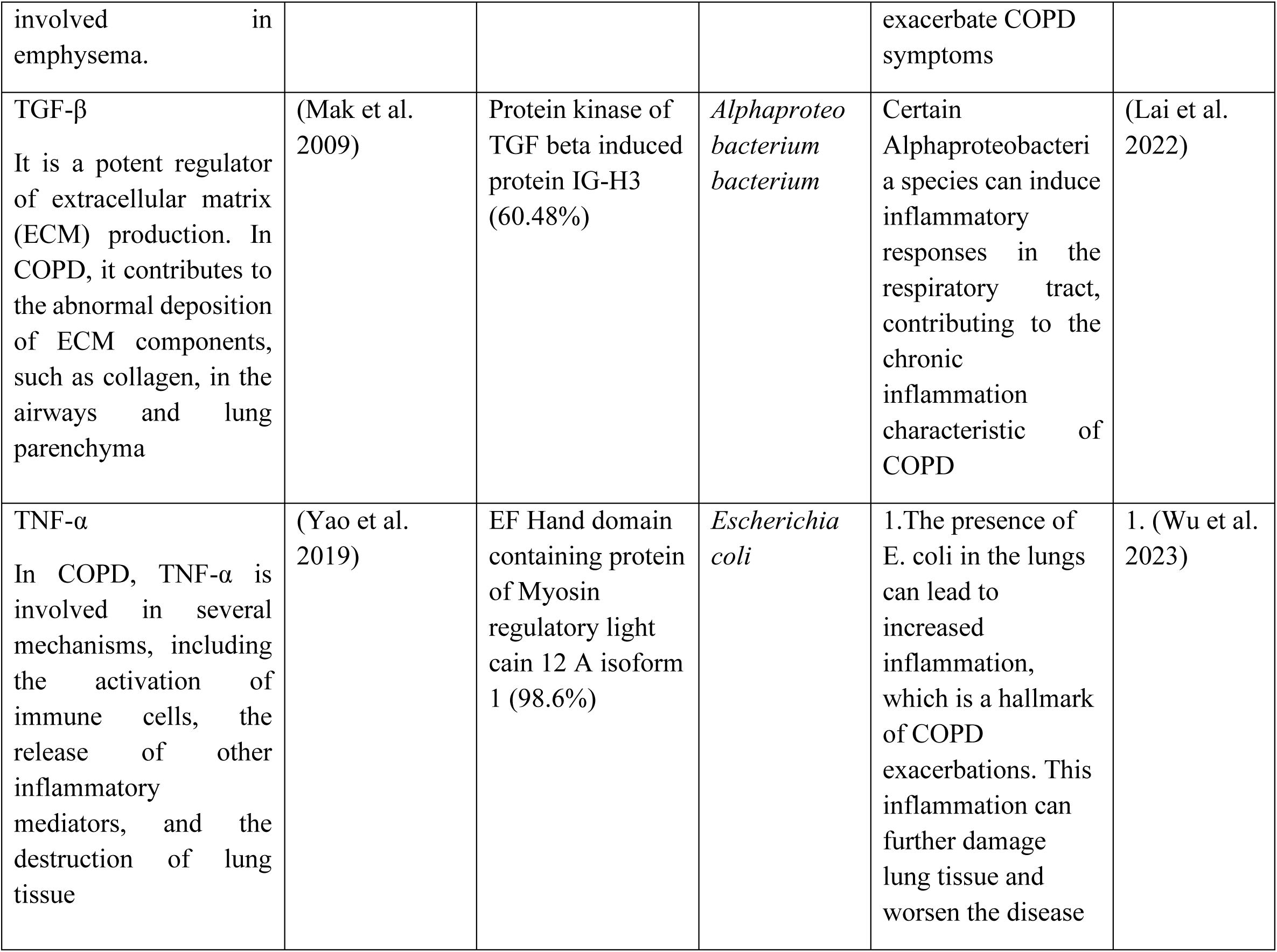
COPD linked human proteins, their known role, microbial homologues along with the name of the microorganism and related literature linking them to the disease.

In COPD, several proteins play a role in triggering the inflammatory response that is responsible for the onset of the disease (Christopoulou et al. 2023; Mak et al. 2009; Yao et al. 2019; Para et al. 2025). It is now known that not only physical factors but also microbial factors like microbiome in human body have a role in progression of the disease. Prominent human proteins responsible for the disease are MMP 1 (Matrix metalloproteinase 1), MMP 2 (Matrix metalloproteinase 2), MMP 14 (Matrix metalloproteinase 14), TNF Alpha (Tumor necrosis factor alpha), TGF Beta (Tumor growth factor Beta). These above proteins have a very important role in COPD disease(Liao et al. 2017; Li et al. 2014). MMP 1 is a type of immune protein which primarily targets type I, II, III,VII collagens that are essential components of the ECM in the lung tissue. In COPD, an imbalance between matrix-degrading enzymes (like MMP-1) and tissue inhibitors of metalloproteinases (TIMPs) leads to excessive degradation of the lung’s structural proteins. This results in the destruction of the alveolar walls, a hallmark of COPD. It play crucial role in extracellular matrix (ECM) degradation and airway remodelling in COPD. It is involved in NF Kappa Beta (NF–kB) and MAPK signalling pathways which regulate inflammation and tissue destruction(Li et al. 2014). MMP 2 is a type of immune protein that degrade various collagens and other ECM proteins (including elastin and fibronectin), can lead to the breakdown of lung architecture during COPD. It play crucial role in extracellular matrix remodelling and airway tissue degradation in COPD. It is involved in TGF Beta/ SMAD pathway, which regulate fibrosis and airway remodelling, MMP 2 Contribute to NF kB mediated inflammation, influencing immune response in COPD. MMP 12 play an important role in elastin degradation and extracellular matrix remodelling in COPD. Elastin is a key component of the lung’s elastic tissue, which provides the lung with its ability to expand and contract. MMP-12 breaks down elastin, contributing to the loss of elasticity in the lungs, a hallmark of emphysema, a common feature of COPD. It is involved in NF kB pathway and MAPK signalling pathway which regulate inflammation and tissue destruction in COPD(Wu et al. 2012). Researchers found that higher amount of COPD patients express higher level of MMP 12 in bronchoalveolar lavage fluid, contributing to excessive extracellular matrix degradation. MMP 14 is a type of immune protein which is involved in emphysema. The damage of the lung alveoli (tiny air sac) is called emphysema(Li et al. 2009). It involves in TGF Beta / SMAD signalling pathway which regulate fibrosis and airway remodelling. TGF Beta is an immune protein which contribute to the abnormal deposition of ECM components such as collagen in the airways and lung parenchyma. TGF Beta contribute to fibrotic epithelial mesenchymal transition (EMT) in COPD patients.It involves in SMAD signalling. TNF Alpha is a type of immune protein in COPD patients. In COPD, TNF-α is involved in several mechanisms, including the activation of immune cells, the release of other inflammatory mediators, and the destruction of lung tissue. It is involved in MAPK signalling pathway which contribute to disease progression by promoting inflammation and tissue damage. This are the reasons why I choose this proteins for in silico evaluation. These all above human proteins (MMP 1, MMP 2, MMP 14, TGF Beta, TNF Alpha) are BLAST in NCBI database against selecting bacteria organism because as human contain microbiome inside (Christopoulou et al. 2023; Yao et al. 2019). The blast results obtained (Table 1) show the homologous bacterial protein and the bacteria contain the homologous protein of that human protein. The human protein MMP 1 show homology with Matrix family metalloprotease of Chain A Interstitial Collagenase protein with 54.88% percentage identity (Homology) and this protein can be found in *Bacillus thuringiensis* bacteria. It has been reported by (Barfod et al. 2010) that exposure to Bt biopesticides can induce sub-chronic lung inflammation. This inflammation could potentially contribute to the development of chronic lung diseases,including COPD. The human protein MMP 2 show homology with Sugar porter family MFS transporter of solute carrier family 2 protein of *Bifidobactrium samirii* with 34.73 % homology. It has been reported by (Lai et al. 2022) that some Bifidobacterium species are known for their anti-inflammatory properties, which could potentially help in reducing lung inflammation associated with COPD. The human protein MMP 14 show homology with Apolipoprotein A1/A4/E family protein of Chain C Apolipoprotein A-I protein of *Klebsiella pneumonia* bacteria with 67.39 % homology. It has been reported by (Janssen Axel et al. 2023) that *K. pneumoniae* can colonize the respiratory tract and lead to infections, particularly in individuals with compromised lung function, such as those with COPD. This colonization can trigger inflammatory responses and exacerbate COPD symptoms. The human protein TGF Beta show homology with Protein kinase of TGF beta induced protein IG-H3 of Alphaproteobacterium with 60.48 % homology. It has been reported by (Lai et al. 2022) that Certain Alphaproteobacteria species can induce inflammatory responses in the respiratory tract, contributing to the chronic inflammation characteristic of COPD. The human protein TNF Alpha show homology with EF Hand domain containing protein of Myosin regulatory light chain 12 A isoform 1 protein of *Escherichia coli* with 98.6 % homology. It has been reported by (Wu et al. 2023) that The presence of E. coli in the lungs can lead to increased inflammation, which is a hallmark of COPD exacerbations. This inflammation can further damage lung tissue and worsen the disease. Next I did molecular docking of this all human proteins with herbal drugs and the homologous microbial protein with the same herbal drugs to see the interaction.

The phylogenetic tree analysis (data not shown) of the homologous protein containing protein demonstrates the evolutionary relationships among five bacterial species harboring proteins homologous to COPD associated human proteins. *Escherichia coli* (WP_248798910.1) appears as the most distantly related species, branching early with a significantly evolutionary distance (42.76), and contain EF hand domain protein highly homologous (98.6%) to TNF linked Myosin regulatory light chain 12A, *Bacillus thuringiensis* (WP_255299030.1), which branches next, contain a metalloprotease from the matrix family with 54.88% homology to MMP 1 suggesting a conserved proteolytic function. Alphaproteobacterium (MDA8032410.1) shares a closer ancestry with *Bacillus thuringiensis* and carries a kinase protein related (60.48%) to TGF Beta signaling, indicating its potential role in immune modulation. Next analysis is the bacteria *Bifidobacterium samirii* (WP_338142198.1) and *Klebsiella pneumonia* (MCP6710965.1) form a distinct clade, implying a closer evolutionary link and functional similarity. *Bifidobacterium samirii* contain a sugar porter family MFS transporter homologous (34.73%) to the solute carrier family, analogous to MMP 2’s role in extracellular matrix (ECM) regulation. *K. pneumonia* possess an apolipoprotein A1/A4/E family protein showing 67.39% similarity to MMP 14, suggesting potential involvement in lipid metabolism and inflammation. The tree topology supported by a strong bootstrap value of 100, underscores the robustness of these ancestral grouping (data not shown). This evolutionary perspective suggests that microbial proteins structurally and functionally similar to COPD related human proteins could influence host pathways. Such findings provide a foundation for understanding microbial contribution to COPD pathogenesis through molecular mimicry or pathway modulation. Overall, the phylogenetic tree analysis highlights the importance of conserved microbial proteins in the host microbiome interface in COPD.

## Conclusion and Future Prospect

This study highlights the potential role of microbial proteins—structurally similar to human inflammatory proteins—in influencing COPD pathophysiology. In silico analyses revealed that several herbal compounds, including menthol, gingerol, and oleanolic acid, exhibit promising binding interactions with both microbial and human proteins implicated in COPD. These findings suggest that certain herbal agents may possess anti-inflammatory and antioxidant properties beneficial for symptom management. Additionally, our results reinforce the emerging understanding of the gut–lung axis and the broader impact of the microbiome on COPD development.

However, most current evidence remains theoretical or preclinical, with limited clinical validation. The therapeutic potential of herbal medicines and microbiome-modulating strategies in COPD requires rigorous in vivo studies and well-designed clinical trials to assess efficacy, safety, dosing, and long-term outcomes. Future research should focus on unraveling the complex host–microbiome interactions and determining how microbial shifts influence treatment responses. Integrating herbal pharmacology with microbiome science may ultimately pave the way for innovative, natural, and personalized therapeutic options for COPD.

## Supporting information

Sipplemental Table 1

## Notes

### Competing Interest Statement

The authors have declared no competing interest.

