## Supplementary material for "*In-Silico* Investigation Reveals a Potential Functional Role of Human Microbiome in Chronic Obstructive Pulmonary Disease": Sipplemental Table 1

**Snapshot of Docking Results**

| **Docking Datatable of Human protein with Oleanolic acid** | **Docking Datatable of Microbial protein (Homologous of human protein) with Oleanolic acid** |
| --- | --- |
| A.  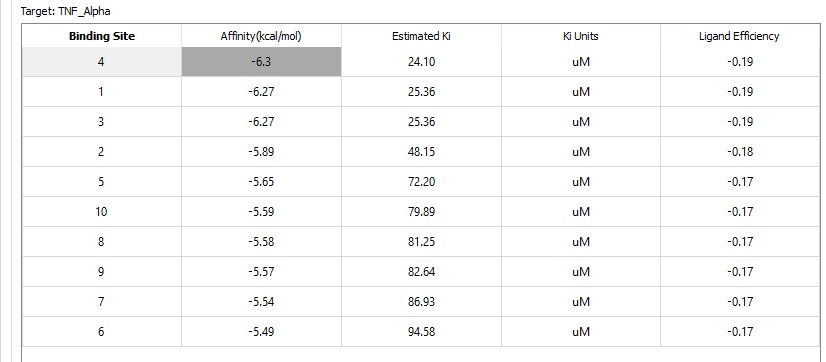 | A’.  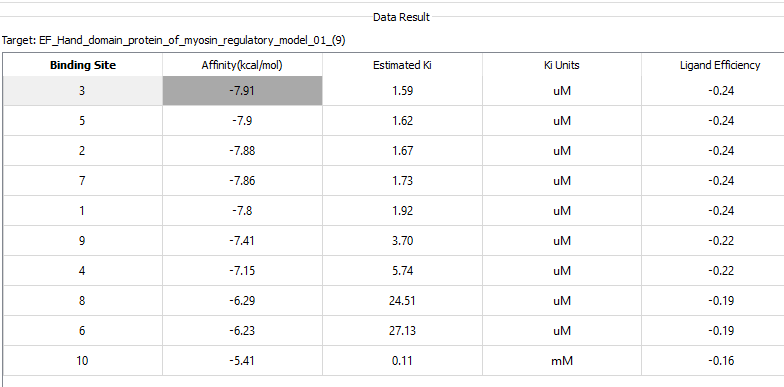 |
| B.  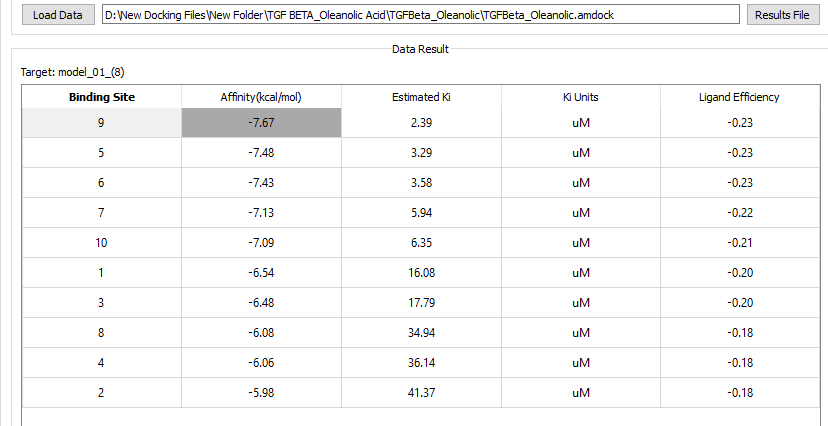 | B’.  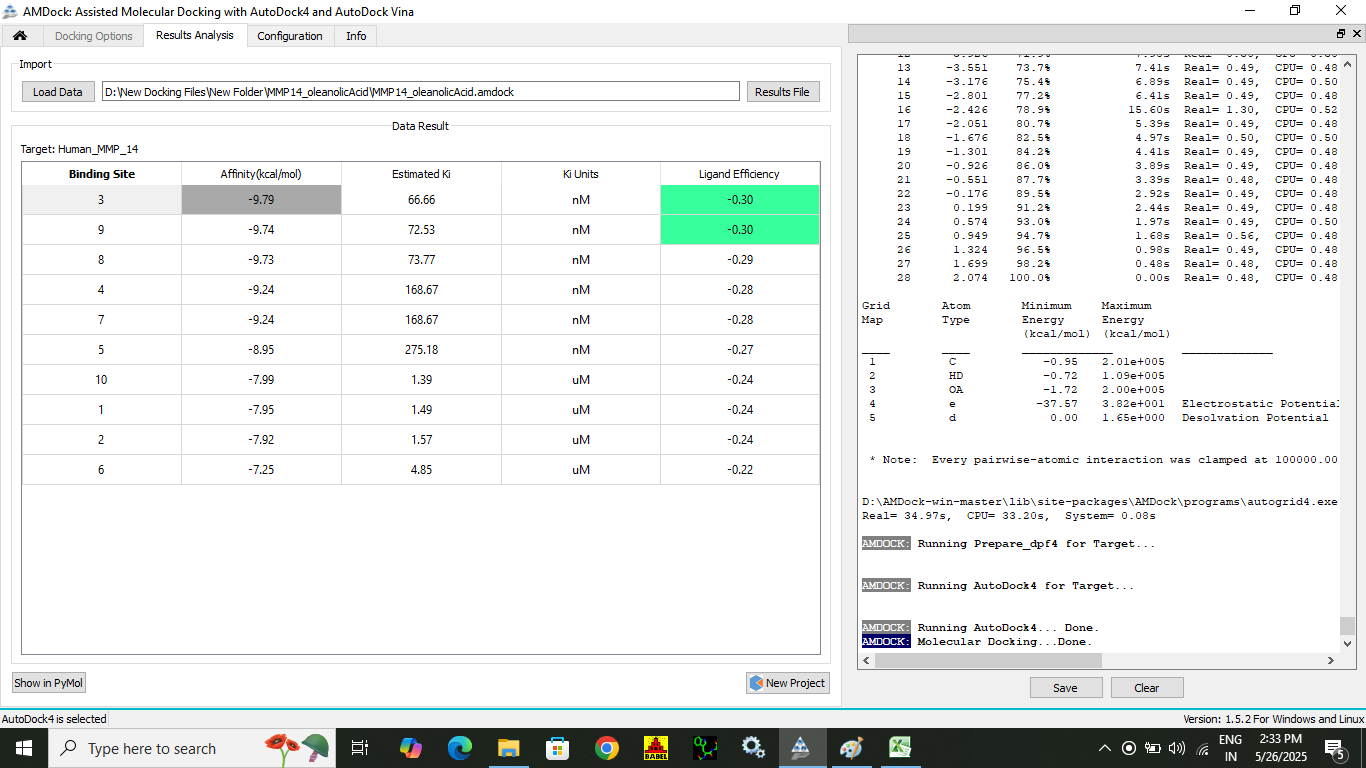 |
| C.  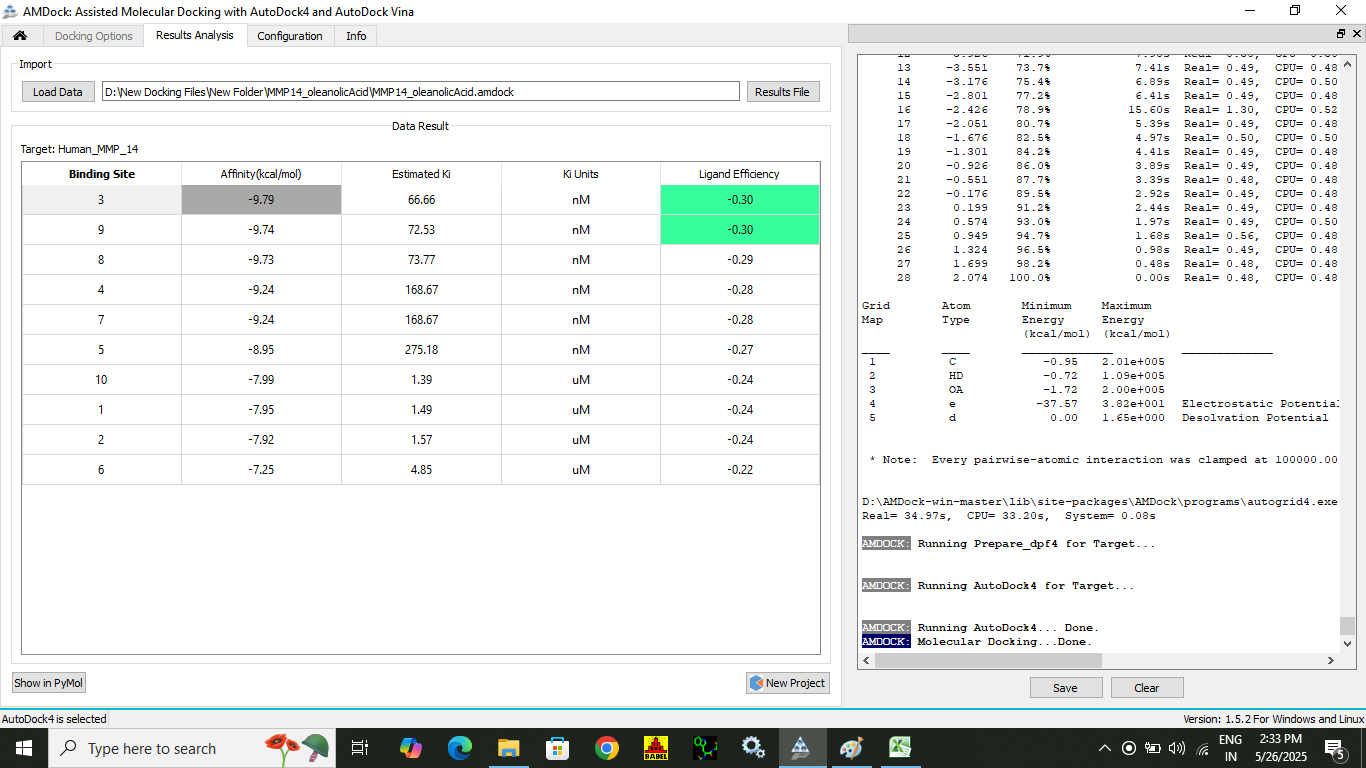 | C’.  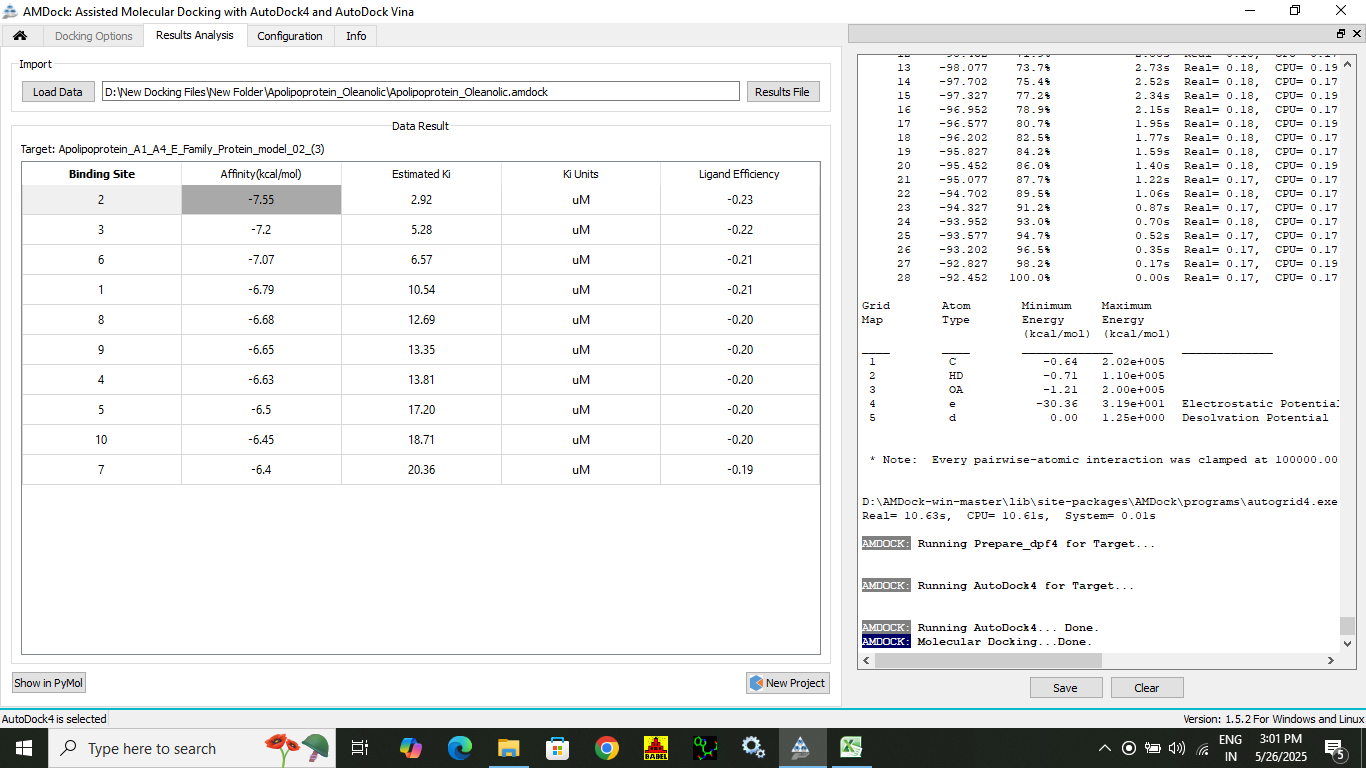 |
| D.  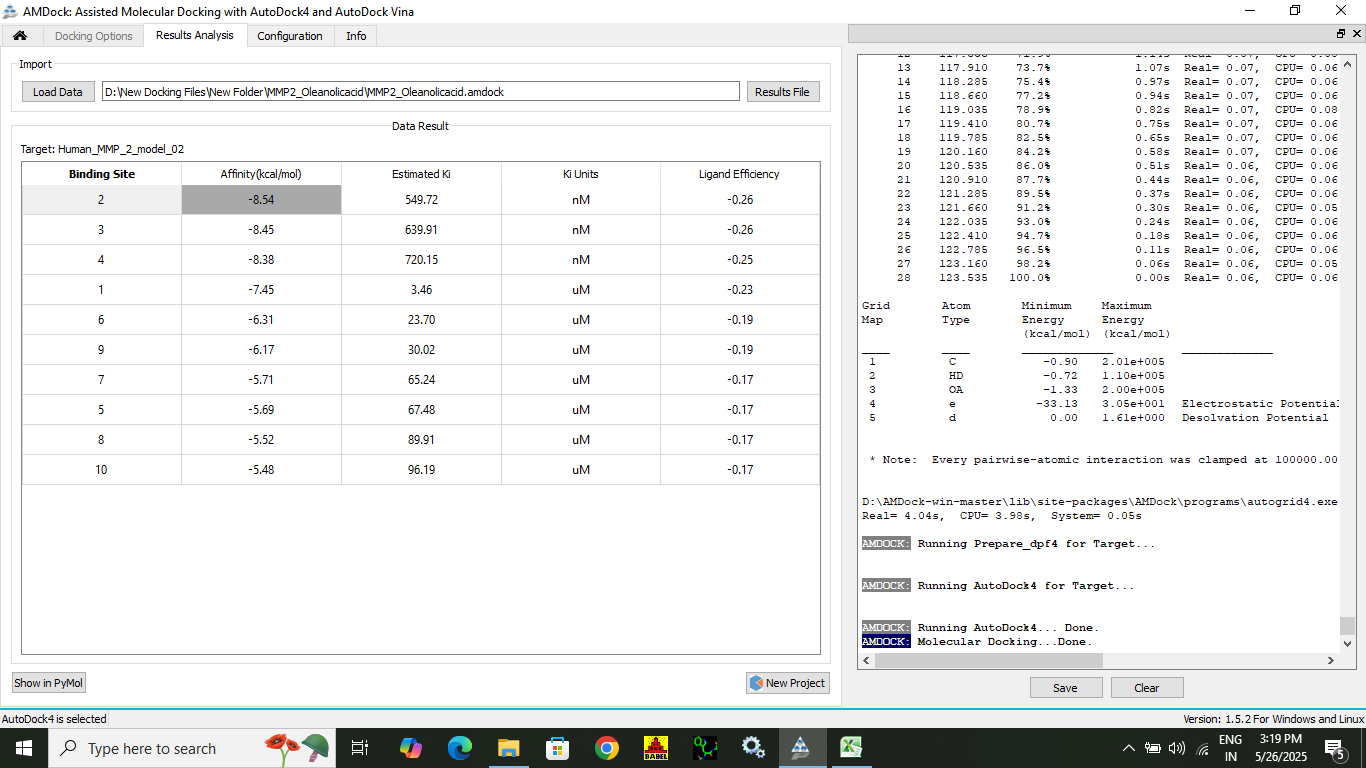 | D’.  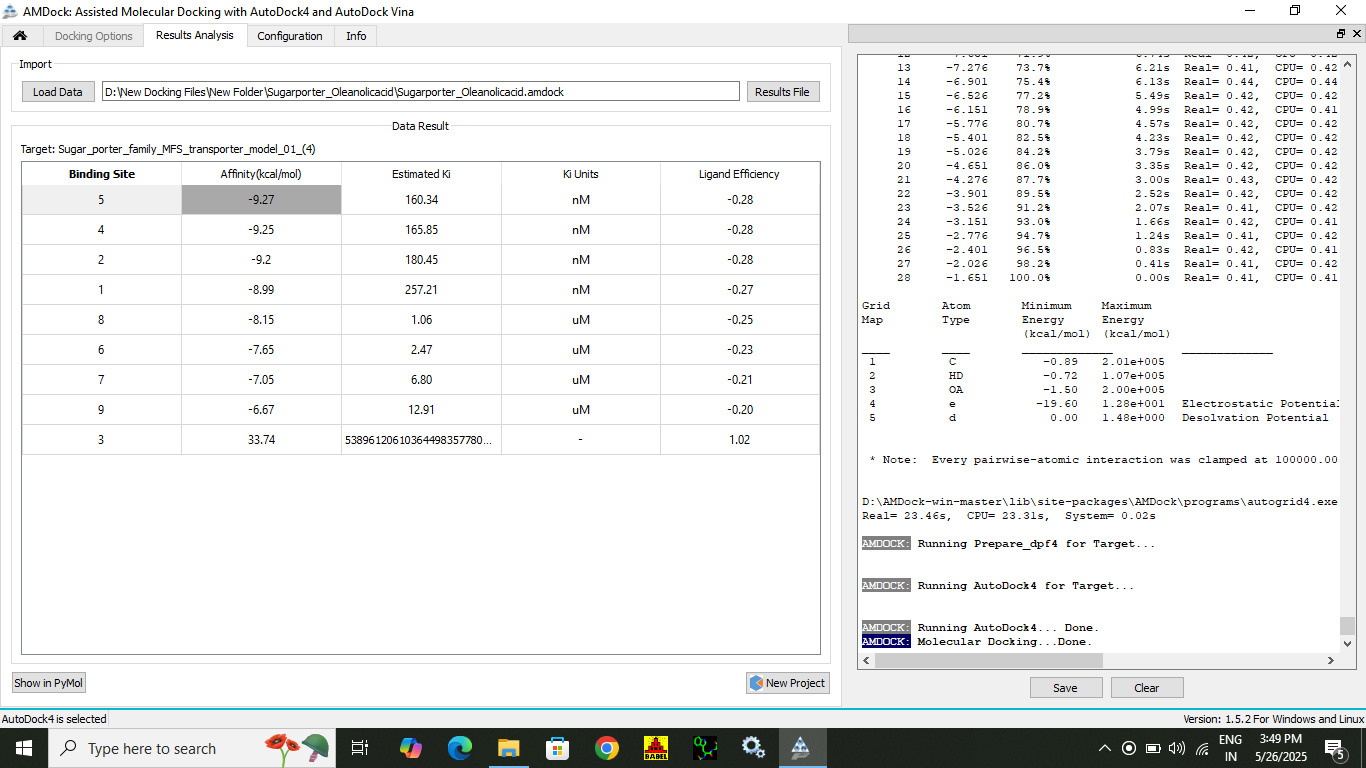 |
| E.  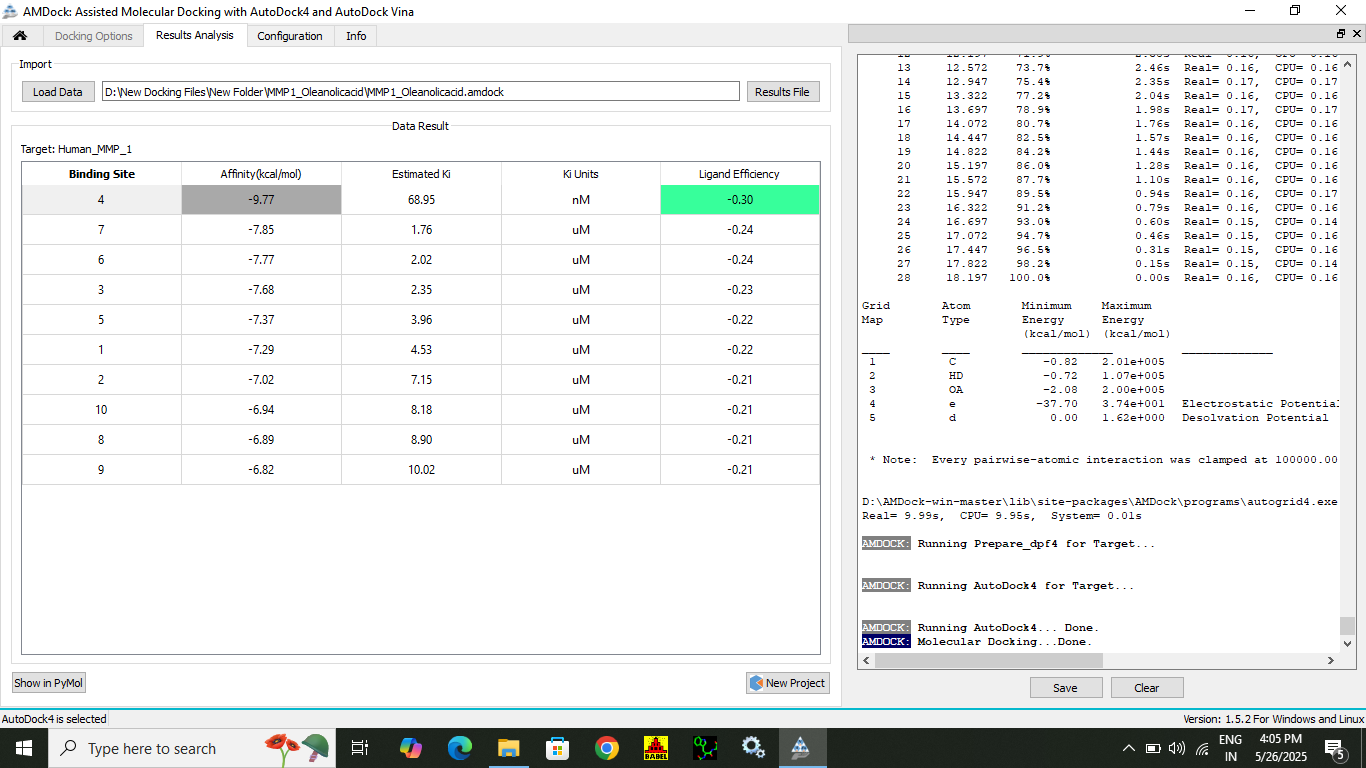 | E’.  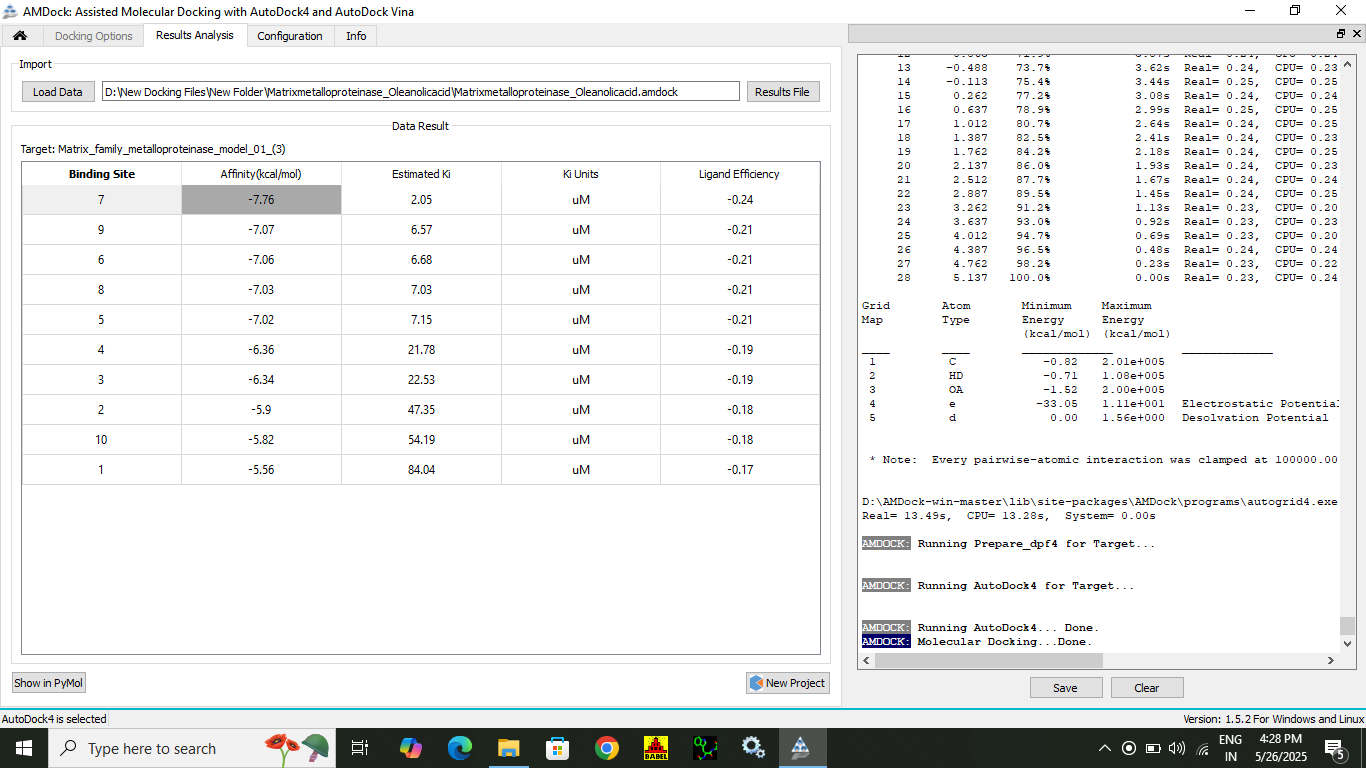 |
